# PIEZO1-Mediated Mechanotransduction Regulates Collagen Synthesis on Nanostructured 2D and 3D Models of Fibrosis

**DOI:** 10.1101/2024.06.30.601386

**Authors:** Neda Rashidi, Natalia S. Harasymowicz, Alireza Savadipour, Nancy Steward, Ruhang Tang, Sara Oswald, Farshid Guilak

## Abstract

Progressive fibrosis causes tissue malfunction and organ failure due to the pathologic accumulation of a collagen-rich extracellular matrix. In vitro models provide useful tools for deconstructing the roles of specific biomechanical or biological mechanisms involved in these processes and identifying potential therapeutic targets. In particular, recent studies have implicated cellular mechanosensing of substrate micro- and nanoscale architecture as a regulator of fibrosis. In this study, we investigated how the mechanosensitive ion channel PIEZO1 and intracellular mechanotransduction pathways influence fibrotic gene and protein expression in adipose-derived stem cells (hASCs). Specifically, we examined the role of PIEZO1 and the mechano-sensitive transcription factors YAP/TAZ in sensing aligned or non-aligned substrate architecture to regulate collagen formation. We utilized both 2D microphotopatterned substrates and 3D electrospun polycaprolactone (PCL) substrates to study the role of culture dimensionality. We found that PIEZO1 regulates collagen production in hASCs in a manner that is sensitive to substrate architecture. Activation of PIEZO1 induced significant morphological changes in hASCs, particularly when they were cultured on aligned substrates. While YAP translocated to the cytoplasm following PIEZO1 activation, depleting YAP and TAZ did not change collagen expression significantly downstream of PIEZO1 activation, implying that YAP/TAZ translocation out of the nucleus and increased collagen production may be independent outputs of PIEZO1 activation. Our studies demonstrate a role for PIEZO1 in cellular mechanosensing of substrate architecture and provide targetable pathways for treating fibrosis as well as for enhancing tissue-engineered and regenerative approaches for fibrous tissue repair.

## Introduction

Tissue fibrosis is a pathologic process characterized by the formation of a fibrous, collagen-rich extracellular matrix (ECM), increased tissue stiffness, and a loss of normal capacity to repair tissue damage and restore homeostasis. Particularly, due to increasing rates of obesity worldwide, there has been growing interest in obesity-related pathologies such as adipose tissue fibrosis [1]. Alteration in ECM mechanical properties is believed to alter crucial signals that regulate cell function, leading to continual progression of this condition [2, 3]. Despite its prevalence across various soft tissues (including adipose tissue, heart, liver, skin, kidney, and eye), effective anti-fibrotic treatments are lacking [4]. Consequently, fibrosis often necessitates organ transplantation, and is a primary cause of morbidity and mortality in a variety of chronic inflammatory disease [5]. As the precise cellular and molecular mechanisms underlying its development, maintenance, and resolution remain incompletely understood, more study is needed [6].

Conventional methods have failed in identifying inhibitors of soluble factor signaling pathways in fibrosis therapy [7], prompting the exploration of mechanobiological pathways [2]. Recognizing the significant role of mechanical forces and tissue stiffness in fibrosis development, recent investigations have delved into mechanotransduction [8-13], informing the development of strategies for modulation or reversal of fibrotic remodeling. In recent research, engineered biomimetic in vitro models have emerged as viable alternative to animal models, seeking to replicate the physical characteristics of in vivo niche [14], aiding in understanding mechanobiological pathways in fibrosis progression [7, 15]. Consideration of dimension, substrate architecture, and stiffness is crucial in disease model construction using in vitro models.

PIEZO1, a mechanosensitive ion channel, is activated when membrane tension increases [16]. It plays crucial roles in human physiology such as touch sensation, and cell mechanosignaling [16-23]. Studies have also shown that PIEZOs are involved in pathology [24-26], including fibrosis [10, 13, 27]. For example, GsMTx4 (a small molecule PIEZO1 inhibitor) can alleviate kidney fibrosis, while Yoda1 (a small molecule PIEZO1 activator) promotes profibrotic responses in renal HK2 cells [9]. Overall, prior studies have identified PIEZO1 as a potential therapeutic target for fibrotic conditions in various organs, including the renal [9, 10, 13] and cardiac fibrosis [28]. In all these cases, PIEZO1 had pro-fibrotic effects. In adipose tissue, PIEZO1 is implicated in adipocyte plasticity, with increased expression in obese mice leading to fibrotic adipose tissue development [27, 29]. While in another study on adipose tissue, PIEZO1 knockdown showed no significant change in fibrotic gene expression, such as collagens, despite excessive collagen synthesis and deposition being hallmark of fibrosis [30]. A recent study also suggested PIEZO1 inhibition as a promising strategy for minimizing fibrosis, especially in organs where fat contributes to fibrotic processes [31]. Thus, further investigation is needed to understand if and how PIEZO1 regulates mechanosensation during the progression of fibrosis in adipose tissue.

YAP (Yes-associated protein) and TAZ (Transcriptional co-activator with PDZ-binding motif), crucial nuclear transducers of mechanical signals [32], have been shown to contribute significantly to the progression of fibrosis [11, 33, 34]. Their regulation and subsequent nuclear translocation are dependent on RhoGTPase activity and actomyosin cytoskeleton tension [35, 36]. In the context of fibrotic diseases, increased expression and nuclear localization of YAP/TAZ have been observed in remodeled fibrotic lungs [11, 12] suggesting their involvement in the fibrotic process. Additionally, ECM stiffening ex vivo and liver damage in vivo have been demonstrated to activate YAP in hepatic stellate cells during liver fibrosis [37]. Notably, targeted inhibition of YAP through knockdown or pharmacological approaches prevents the activation of hepatic stellated cells and subsequent fibrogenesis in mice models [37]. While YAP and TAZ are primarily associated with fibrosis, they also play a role in adipose tissue biology, where TAZ regulates adipogenic activity in mature adipocytes [38], and the knockout of both YAP and TAZ negatively impacts adipose tissue expansion in mice [39]. Together, these findings highlight the role of YAP and TAZ in fibrotic diseases as well as their involvement in adipose tissue remodeling, potentially through their mechanosensitive regulation. However, the precise role of PIEZO1 or YAP/TAZ pathways in tissue fibrosis remains to be determined, and a better understanding could provide important opportunities for identifying new therapeutic targets for this condition.

Cells utilize mechanosensitive ion channels as a means of perceiving their physical environment. Research indicated that substrate properties and architecture influence the activity of these ion channels. For instance, PIEZO1 has been indirectly implicated in detecting the nanoroughness of substrates in interactions between neurons and astrocytes [40], and it has been observed to influence the differentiation of neural progenitor cells based on substrate stiffness [41]. Additionally, PIEZO1 has been identified as a sensor for confinement during cell migration [42]. Despite these advances, the impact of substrate architecture on PIEZO1-mediated mechanotransduction in collagen production is not fully understood.

Adipose-derived stem cells (ASCs) have gained increasing interest as a valuable cell source to study fibrosis and test potential therapeutic interventions, due to their potentail antifibroitic effects. ASCs are multipotent adult stem cells with self-renewal and multilineage potential, and are easily isolated, making them a suitable cell source for *in vitro* studies. Three methods by which ASCs are thought to counteract fibrosis have been proposed: regenerative activity, direct cell-to-cell contact, and paracrine signaling. They interact with a variety of cell types-including immune cells, endothelial cells, and fibroblasts- all significant contributors to fibrosis. Their ability to counteract fibrosis is thought to arise from releasing growth factors that affect the different cell types they interact with [43-47]. For example, due to their paracrine factors, ASCs demonstrated therapeutic promise for liver fibrosis [48-50]. Moreover, ASCs are hypothesized to play a crucial role in the anti-fibrotic effects of autologous fat grafting, a technique showing promise in treating skin fibrosis, particularly through their secreted factors [51].

The goal of this study was to elucidate PIEZO1 mechanosensing mechanisms in hASCs within a stiff microenvironment, and to understand how substrate architecture influences profibrotic signaling. Both 2D model (microphotopatterning) and 3D models (PCL) were utilized to investigate role of culture dimensionality in-vitro. We hypothesized that PIEZO1 signaling is regulated by the alignment or lack thereof of substrate architecture, modulating Ca^2+^ signaling and collagen production, and that this process is regulated downstream via YAP/TAZ signaling. Our findings indicate a critical role for PIEZO1 in regulating collagen expression, influenced by both 2D and 3D matrix architecture. Our findings shed new light on the mechanosensitive processes involved in both physiologic and pathologic fibrous tissue formation.

## Materials and Methods

### Cell culture

Immortalized human adipose stem cells (ASC52telo, ATCC SCRC4000™) were expanded in monolayer in Mesenchymal Stem Cell (MSC) Basal Medium (ATCC, PCS-500-030). 10 mL of MSC supplement and 6 mL of LAlanyl-L-Glutamine (ATCC, PCS-500-040) and 0.2 mg/mL G418 (Sigma, G8168) were included as medium supplements. hASCs were seeded (30,000 cells/cm^2^) onto aligned patterned or unpatterned (flat) surfaces in the 2D model, or for cell culture on PCL mats with aligned or random fibers. Cells were cultivated for 7 days; cultured media (mesenchymal stem cell basal medium with ascorbic acid (50 μg/ml) was replaced every three days.

### Preparation of micropatterned substrates

On large (75.2 × 50.4 × 1 mm) microscope coverslips (Ted Pella, Redding CA), cell-adhesive patterns were produced by photoablating thin (150 nm) polyvinyl alcohol hydrogel films (PVA) (Sigma-Aldrich, St. Louis, MO) utilizing lithography techniques. Briefly, large cover glasses (Ted Pella, Redding, CA) were activated as described in [52], and their surface was then spin-coated with PVA and S1805 photoresist. Coverslips were baked for one minute at 100°C. Either a laser writer or a mask aligner wrote the patterns. The photoresist was subsequently developed with a developer for 1 minute. After 5 minutes of drying at 70°C, cover glass surfaces were etched using reactive ion etching (RIE) device. Photoresist was removed with remover PG (Fisher Scientific, Waltham, MA) for 4 minutes. Cover glasses were then washed with deionized water. UV light was utilized to disinfect cover glasses for 1 hr. Micropatterns comprised of two micro-scale architectures: 1) aligned, with straight parallel lines (2 μm wide, 5 μm center-to-center distance), and 2) unpatterned (flat), with fully ablated areas (2 cm by 0.2 cm), as depicted in (Fig. S2). Patterns were stored at 4°C in 1X PBS (Gibco). This technique produces patterns that are stable for one month prior to cell seeding and two weeks in culture. Pattern area was coated with 20 μg/ml fibronectin (Sigma-Aldrich, St. Louis, MO) to functionalize the ablated cell-adhesive regions prior to cell culture. Then, cover glasses were incubated at 37°C for 1 hr, washed three times with 1X PBS, and seeded with hASCs. The selective adsorption of fibronectin to patterned surfaces was visualized using fluorescently labeled (green) fibronectin (Cytoskeleton, Inc, Denver, CO). Cells began to align in the pattern direction 2 hrs after seeding (Fig. S2).

### Preparation of 3D electrospun polycaprolactone (PCL) substrates

Electrospun polycaprolactone (PCL) substrates with aligned and random fibers (Nanofiber Solutions, Dublin, OH) were used as our 3D cell culture model. 24 well plates or plate inserts were used for all RT-qPCR, immunolabeling and Western blot. They were rinsed 2-3 times with 1x PBS and fibers allowed to air-dry. Mats were coated with 20 μg/ml fibronectin (Sigma-Aldrich, St. Louis, MO) for 2 hrs at 37°C to functionalize fibers prior to cell culture. After 3 washes of 1x PBS, mats were pre-incubated in media for at least 2 hrs at 37°C. The media was removed by aspiration and cells were seeded on substrates with aligned or random fibers.

### Immunostaining

The media was aspirated, and the cells were washed twice with 1x PBS. Cells were fixed with 4% paraformaldehyde (Electron Microscopy Sciences, Hatfield, PA) in 1x PBS at room temperature (RT) for 15 minutes, followed by 3 washes with 1x PBS. hASCs were then permeabilized with 0.1% Triton X-100 (Sigma-Aldrich, St. Louis, MO) in 1x PBS for 5 minutes at room temperature. Samples were then rinsed twice with 1x PBS followed by blocking with goat serum (Vector® laboratories, Newark, CA) for 30 minutes in RT. Primary antibodies for YAP (Cell signaling, Danvers, MA, 1:200), PIEZO1 (Novus Biologicals, Centennial, CO, 1:25), Collagen I (Cell Signaling, Danvers, MA, 1:1000) were use. Samples were covered by primary antibodies and incubated for 1 hr at RT, followed by three washes with 1x PBS tween for 5 minutes each. The secondary antibody was diluted according to manufacturer protocol in 1x PBS tween and incubated at RT for 1 hr. The samples were then washed three times for 5 minutes each with 1x PBS tween and stained for 20 minutes with TRITC-coupled Phalloidin (Invitrogen, Carlsbad, CA). After three washes with 1X PBS, cells were mounted using mounting media with DAPI (Vectashield, Newark, CA) on a microscope slide (Fisher Scientific, Waltham, MA).

### Picrosirius Red Staining

hASCs were fixed for 15 minutes at RT with 4% paraformaldehyde (Electron Microscopy Sciences, Hatfield, PA) in 1x PBS. The cells were rinsed three times with 1x PBS and treated for 1 hour at RT with picrosirius red staining solution (0.1% Direct Red 80, 0.1% Fast Green FCF, dissolved in a saturated aqueous solution of picric acid, which contains 1.2% picric acid in water) followed by incubation with 1% acidified water (acetic acid). After 5 minutes, cells were washed twice with 1x PBS and afterwards kept at 4°C until imaging.

### Real-time PCR

Following the manufacturer’s instructions, RNA was extracted using a total RNA purification kit (NORGEN, Auburn, WA) followed by cDNA synthesis utilizing iScript cDNA kit (BIORAD, Hercules, CA). Quantitative real time-PCR using iTaq Universal SYBR GREEN Supermix (BIORAD, Hercules, CA) was used to assess the mRNA expression level of the target genes.

### Primer Design

Using the NCBI Primer-BLAST and Primer3 websites, primers for the target genes were designed and purchased from idtDNA. Primers are listed in Table S1. Using the 2(-ΔΔCT) approach, the relative fold change was assessed for gene expression analysis. The expression of the target gene was normalized relative to the housekeeping gene (GAPDH) and the gene expression of the control (aligned) samples.

### siRNA knockdown

Small interfering RNA (siRNA) transfections were performed utilizing Dharmafect1 (Horizon) according to the manufacturer’s guidelines. On-TARGET plus siRNA pools were used to transfect target genes. The sequences of siRNA pool for each gene are listed in Table S2. The total concentration of all siRNAs employed for transfection was 25 nM. 48 hours after transfection, RNA samples were isolated, and 96 hours after transfection, protein samples were isolated. siRNA method was used to knock down *PIEZO1*, *YAP* and *TAZ* genes on our hASCs culture both on micropatterns and PCL substrates.

### Western blotting

Cells were lysed in RIPA buffer (Cell Signaling, Danvers, MA) with 2.5% CHAPS (Sigma-Aldrich, St. Louis, MO) and protease inhibitor (ThermoFisher, St. Louis, MO). Then, protein content was assessed using the BCA Assay (Pierce). 25 µg of proteins were separated in each well using 6% sodium dodecyl sulfate-polyacrylamide gel electrophoresis gel with pre-stained molecular weight markers (ThermoFisher St. Louis, MO) and transferred to PVDF (polyvinylidene) membranes. The PVDF membranes were incubated overnight at 4°C with anti-PIEZO1 (Protein tech, Rosemont, IL) and anti-GAPDH (Protein tech, Rosemont, IL) antibodies. The relevant HRP-conjugated secondary antibodies (Cell Signaling, Danvers, MA) were utilized for detection. Using the iBright FL1000 Imaging System, immunoblots were imaged and evaluated (Thermo Fisher, St. Louis, MO). After normalization with the signal intensity of GAPDH, histograms reflect the signal intensity and area of protein bands in arbitrary units.

### Imaging with Confocal microscopy

hASCs cultured on fibronectin coated micropatterns or PLC mats were imaged using a confocal microscope (LSM 880, Zeiss, Dublin, CA). Z-stack images were acquired with a 40x objective. Images were taken from at least four field of views from each sample. Alexafluor 647 was used for secondary antibody YAP, Alexafluor 488 was used for phalloidin, 405 fluorophore was used for DAPI.

### Ca^2+^ Imaging

Cells were cultivated in MatTek Petri dishes coated with fibronectin. Before imaging, hASCs were stained for 1 hr with Fura red-AM (ThermoFisher Scientific, St. Louis, MO) and Fluo-4-AM (ThermoFisher Scientific, St. Louis, MO). Before imaging, the staining medium was removed and replaced with imaging media (HBSS media). The images were captured with a confocal microscope (LSM 880, Zeiss, Dublin, CA) fitted with a 10x objective. During imaging, the temperature was maintained at 37°C using an incubation stage. One minute of baseline time series photos were captured before the DMSO (control) solution was added to the well. The imaging continued for an additional 2 minutes. The PIEZO1 agonist Yoda1 was added to each well at concentrations of 1 μM, 2.5 μM, 5 μM, or 10 μM for an additional 2 minutes, and the Ca^2+^ signaling response of hASCs was recorded.

### Image Analysis

To quantify the ratio of nuclear/cytoplasmic YAP fluorescence intensity values, we used the following formula: 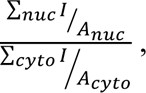 Where ∑*_nuc_ I* and ∑*_cyto_ I* indicate the cumulative intensity values of pixels in nuclear and cytoplasmic region respectively, and *A_nuc_* and *A_cyto_* are the area of the corresponding regions quantified using ImageJ. For the quantification of cell morphology, we employed ImageJ to calculate cell, nucleus area and aspect ratio. Cell aspect ratio was calculated by fitting an ellipse to the cell body or nucleus based on the fluorescent signals from the phalloidin and DAPI channels. The aspect ratio of cell or nucleus was determined as the ratio between the longest and shortest diameters of the fitted ellipse. In Ca^2+^ signaling experiments, a ratiometric study using ImageJ was conducted. After thresholding and using the analysis particle tool, cell pixel intensities were quantified and normalized to control section values. The highest normalized intensity (Fmax/F) represented cell signal. Cells with intensity exceeding the control mean plus five standard deviations were considered to be responding cells. Calculations were performed using a custom MATLAB program.

### Statistical Analysis

For the examination of gene expression, YAP localization and morphological data, a two-way ANOVA followed by a post-hoc test was utilized.

## Results

### hASCs express functional PIEZO1 channels

We first examined the presence and functionality of PIEZO1 in hASCs. By utilizing Western blotting, we confirmed the presence of PIEZO1 protein expression in hASCs (Fig. 1A). When exposed to the PIEZO1-specific chemical agonist Yoda1, hASCs responded by increasing the intracellular Ca^2+^ levels as measured by peak Ca^2+^ signal intensity values (Fig. 1B). This effect was shown to be dose-dependent, 94.4% 10 μM and 83.7% response with 2.5 μM Yoda1 (Fig. 1C). PIEZO1 activation with low doses of Yoda1 (1 μM) showed delayed sustained peak while higher doses of Yoda1 showed shorter peaks with higher signaling (Fig. 1B). These data indicate that ASCs express functional PIEZO1 ion channels.

**Fig. 1.**
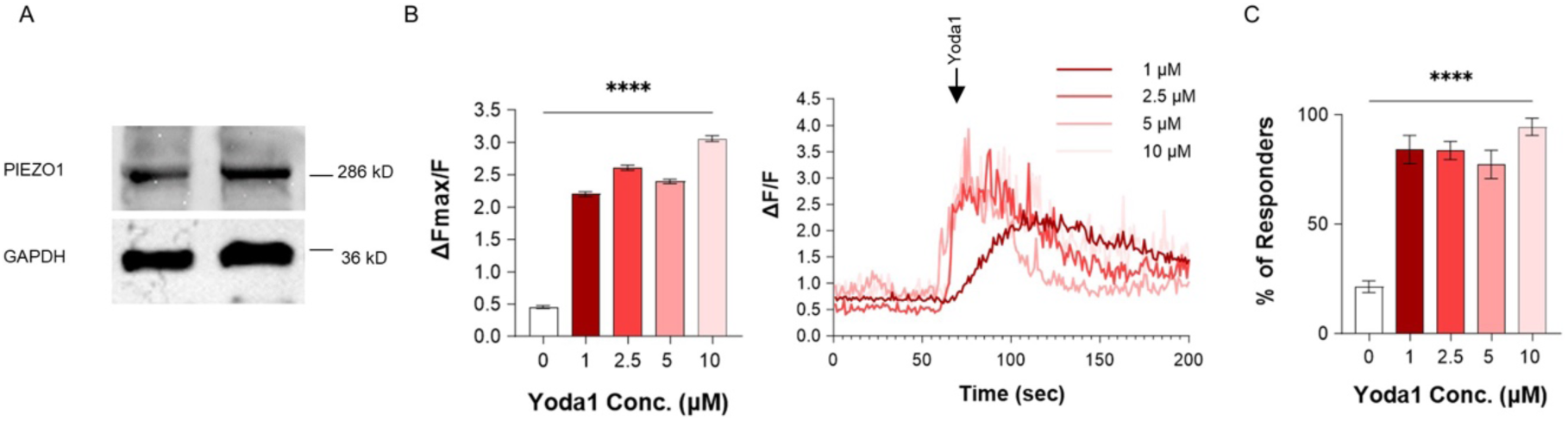
hASCs express functional PIEZO1. A) Protein expression of PIEZO1 in hASCs by western blotting. 286 kD, and 36 kD are expected sizes for PIEZO1, and GAPDH proteins respectively. C) Normalized intracellular Ca^2+^ fluorescence intensity ΔF_max_/F in response to different doses of Yoda1. D) Percentage of cells with Ca^2+^ response for different Yoda1 concentrations. Data presented as mean ± SEM. For C and D, n=3 samples; for group comparison C, D, one-way ANOVA with Dunn post hoc test, \*\*\*\**P<*0.0001.

To directly test the role of PIEZO1 in hASC responses, we used siRNA knockdown [53]. We verified knockdown efficiency in mRNA and protein levels using real time PCR, western blot analysis and immunostaining (Fig S1. A, B, C). Gene expression analysis showed 92.5 % knockdown efficiency in mRNA level and Western blot data showed 73% knockdown efficiency in protein level.

### PIEZO1 modulates collagen production by hASCs

We utilized our multicellular micropattern model system with two configurations: aligned, with straight parallel lines, and unpatterned (flat), with fully ablated areas on glass surface to assess hASCs’ collagen production in response to PIEZO1 modulation on aligned versus non-aligned substrate architecture. As excessive collagen deposition is a hallmark of fibrosis, there has been increasing interest in studying the regulation of collagen expression to better understand the progression of fibrosis [2]. We evaluated collagen synthesis by hASCs cultured on micropatterns using three different conditions: a control group (no treatment), a PIEZO1 antagonist group (treated with 2 µM GsMTx4, with media changes every three days), and a PIEZO1 agonist group (exposed to 2.5 µM Yoda1 for 30 minutes daily). After seven days, we performed Picrosirius red (PSR) staining of cells to visualize collagens. We found that GsMTx4 resulted in increased collagen production, whereas Yoda1 reduced collagen production (Fig. 2A). Moreover, PSR staining of hASCs cultured for seven days on both aligned and flat substrates showed enhanced collagen production in PIEZO1 siRNA-treated groups compared to those treated with non-targeting siRNA (Fig. 2B).

**Fig. 2.**
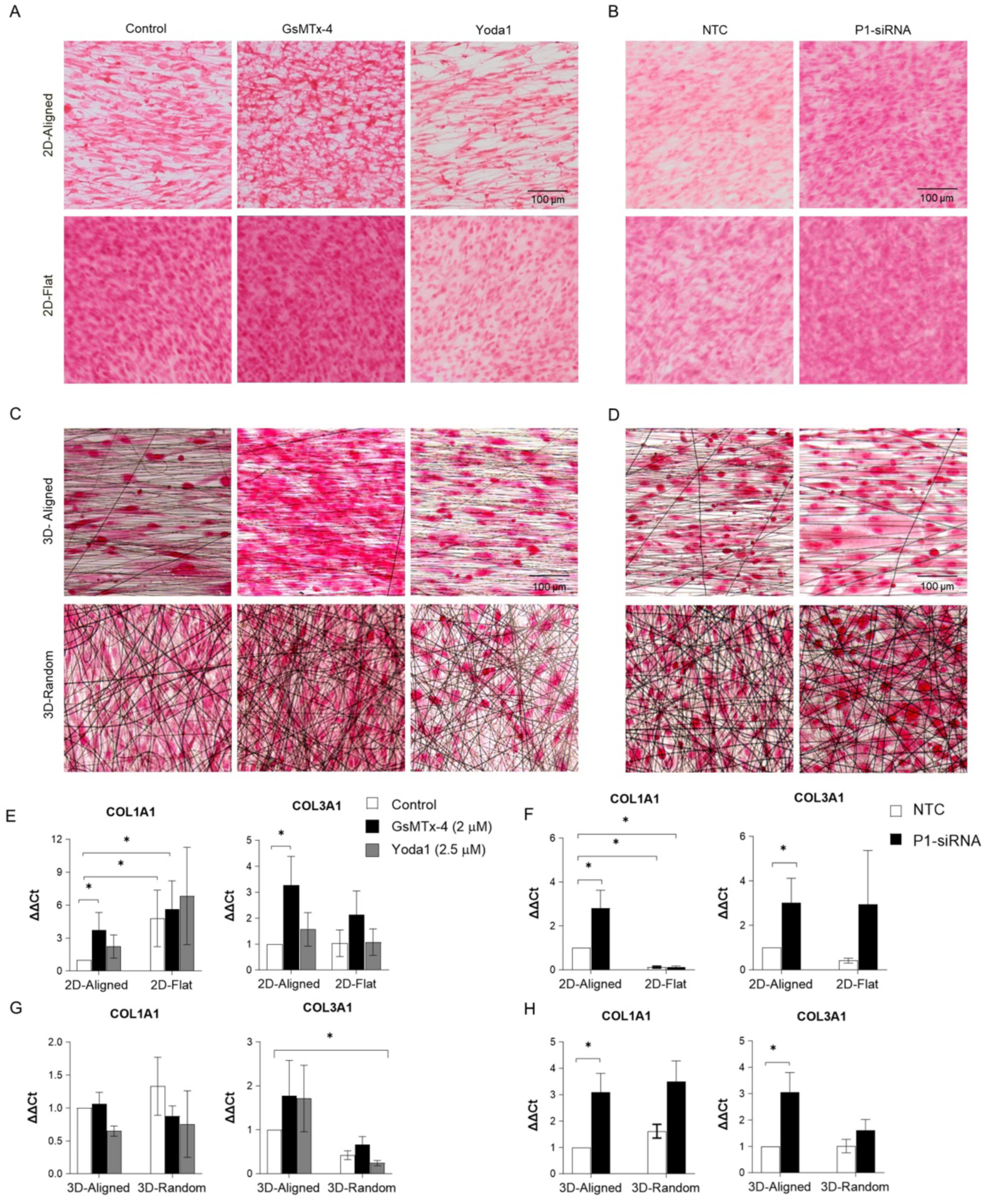
PIEZO1 modulates collagen production in hASCs. A) Picrosirius red staining of hASCs in response to PIEZO1 antagonist (2 μM, GsMTx-4), and PIEZO1 agonist (2.5 μM, Yoda1) in micropatterns. B) Picrosirius red staining of Nontargeting siRNA and PIEZO1 siRNA on micropatterns. C) Picrosirius red staining of hASCs in response to PIEZO1 antagonist (2 μM, GsMTx-4), and PIEZO1 agonist (2.5 μM, Yoda1) on PCL mats. D) Picrosirius red staining of Nontargeting siRNA and PIEZO1 siRNA on PCL mats. E) mRNA levels of *COL1A1* and *COL3A1* normalized to *GAPDH* in response to 2 μM GsMTx-4, and 2.5 μM Yoda on micropatterns. F) mRNA levels of *COL1A1* and *COL3A1* normalized to *GAPDH* in response to Nontargeting siRNA and *PIEZO1* siRNA G) mRNA levels of *COL1A1* and *COL3A1* normalized to *GAPDH* in response to 2 μM GsMTx-4, and 2.5 μM Yoda on PCL mats. F) mRNA levels of *COL1A1* and *COL3A1* normalized to *GAPDH* in response to Nontargeting siRNA and *PIEZO1* siRNA on PCL mats. Data presented as mean ± SEM. For E, n=3 samples; for F, G and H, n= 5 or 6 samples; for group comparison C, D, two-way ANOVA with Dunn post hoc test, \**P<*0.05.

While 2D models provide reductionist systems to test specific architectural features on cell responses, 3D models can provide additional information and better replicate the complex architecture and cellular interactions found in living tissues compared to 2D models. Therefore, collagen production was also investigated in a three-dimensional (3D) model. hASCs were cultured on 3D polycaprolactone (PCL) substrates with aligned and random oriented fibers for a period of seven days. Picrosirius Red (PSR) staining (Fig. 2C) showed that the inhibition of PIEZO1 via GsMTx4 increased collagen production, while activation of PIEZO1 through Yoda1 exposure either reduced collagen synthesis (on 3D-aligned substrates) or had no effect (on 3D-random substrates). In line with our findings in 2D, PIEZO1 knockdown increased collagen production in 3D (Fig. D).

Inhibition of PIEZO1 also led to an upregulation of the collagen genes *COL1A1* and *COL3A1* compared to untreated controls on 2D-aligned surfaces (Fig. 2E). While overall *COL1A1* expression was higher in cells cultured on unpatterned (flat) surfaces, compared to those grown on aligned surfaces, the increase in COL1A1 and COL3A1 gene expression due to PIEZO1 inhibition was not significant on flat surfaces. Treatment with Yoda1 did not have significant effect on collagen gene expressions. Additionally, PIEZO1 knockdown significantly increased *COL3A1* expression in cells on flat substrates and increased both *COL1A1* and *COL3A1* expression in cells on aligned substrates (Fig. 2F).

*COL1A1* and *COL3A1* mRNA levels remained relatively unchanged under exposure to either GsMTx4 or Yoda1 in 3D (Fig. 2G). The only significant difference was in *COL3A1* levels, between cells exposed to Yoda1 and grown on randomly oriented fibers, and the aligned controls (Fig. 2G). However, mRNA levels of both *COL1A1* and *COL3A1* showed a significant, 2.5-3-fold increase following PIEZO1 knockdown in cells grown on aligned 3D substrates (Fig. 2H). Cells on 3D-random substrates did not show a significant change in collagen gene expression. To specifically quantify collagen I protein expression in a 3D context, we used immunolabeling and Western blotting on hASCs cultured on PCL mats. hASCs were exposed to Yoda1 to activate PIEZO1, or *PIEZO1* was knocked down and collagen protein were measured when in the presence or absence of PIEZO1. Western blots showed that PIEZO1 activation was associated with an approximately 2-fold reduction in collagen levels in cells cultured on both aligned and random fibers (Fig. 3A, C). Immunostaining results were consistent with Western blots. Conversely, the knockdown of *PIEZO1* increased collagen at protein levels on both aligned and random fibers (Fig. 3B, D). Taken together, these results indicate that PIEZO1 senses substrate architecture to regulate collagen expressions in hASCs on stiff substrates and this regulation shows dependency to substrate architecture in mRNA levels.

**Fig. 3.**
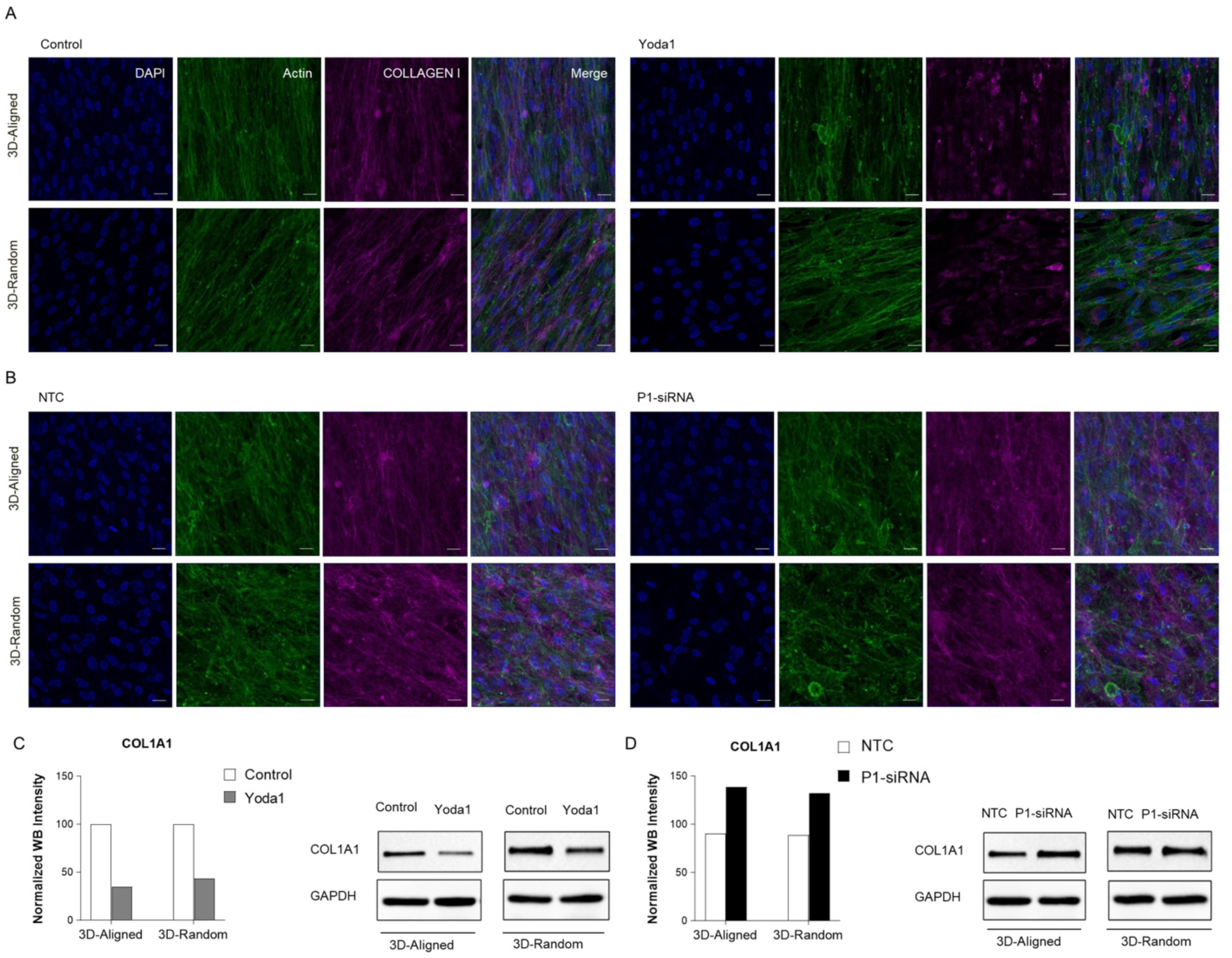
PIEZO1 modulates collagen production in hASCs at the protein level. A) Representative nuclear (blue), actin (green), and Collagen (pink) staining of hASCs cultured on PCL substrates for Control and 2.5 μM Yoda1 groups. B) Representative nuclear (blue), actin (green), and Collagen (pink) staining of hASCs cultured on PCL substrates for Non targeting siRNA and PIEZO1 siRNA groups. C) Protein expressions of Collagen I in control and 2.5 μM Yoda1 groups by western blotting (top). Normalized intensity values of protein expression quantified using WB (bottom). D) Protein expressions of Collagen I in Non targeting siRNA and PIEZO1 siRNA groups by western blotting (top). Normalized intensity values of protein expression quantified using WB (bottom).

### PIEZO1 activation leads to increase cytoplasmic YAP

YAP localization serves as an indicator of cell responses to mechanical cue in the microenvironment; YAP is in its active form when it is localized in the Nucleus [35]. In order to test if the YAP/TAZ pathway was involved in the PIEZO-mediated changes in collagen in hASC cells, we quantified YAP localization following PIEZO1 activation with Yoda1. Nuclear accumulation of YAP or TAZ has been widely used as a primary indicator of the active YAP/TAZ-associated signaling pathways. Nuclear-to-cytoplasmic (N/C) ratio of YAP serves as a read out of cellular mechanosensing [35]. hASCs cultured on either aligned or flat micropatterned surfaces were treated with (or without) Yoda1 for 1 hour. Then, immunostaining and confocal imaging were used to measure changes in YAP localization and fluorescence intensity ratios. Image analysis revealed that the activation of PIEZO1 in hASCs induced a decrease in nuclear/cytoplasmic ratio of YAP (Fig. 4A, C) on either aligned and flat substrates. Our immunofluorescence analyses indicated that PIEZO1 activation on hASCs cultured on PCL mats also decrease in nuclear/cytoplasmic ratio of YAP (Fig. 4B, F). These findings show that PIEZO1 modulation has a significant influence on YAP activity in 2D or 3D cellular environments.

**Fig. 4.**
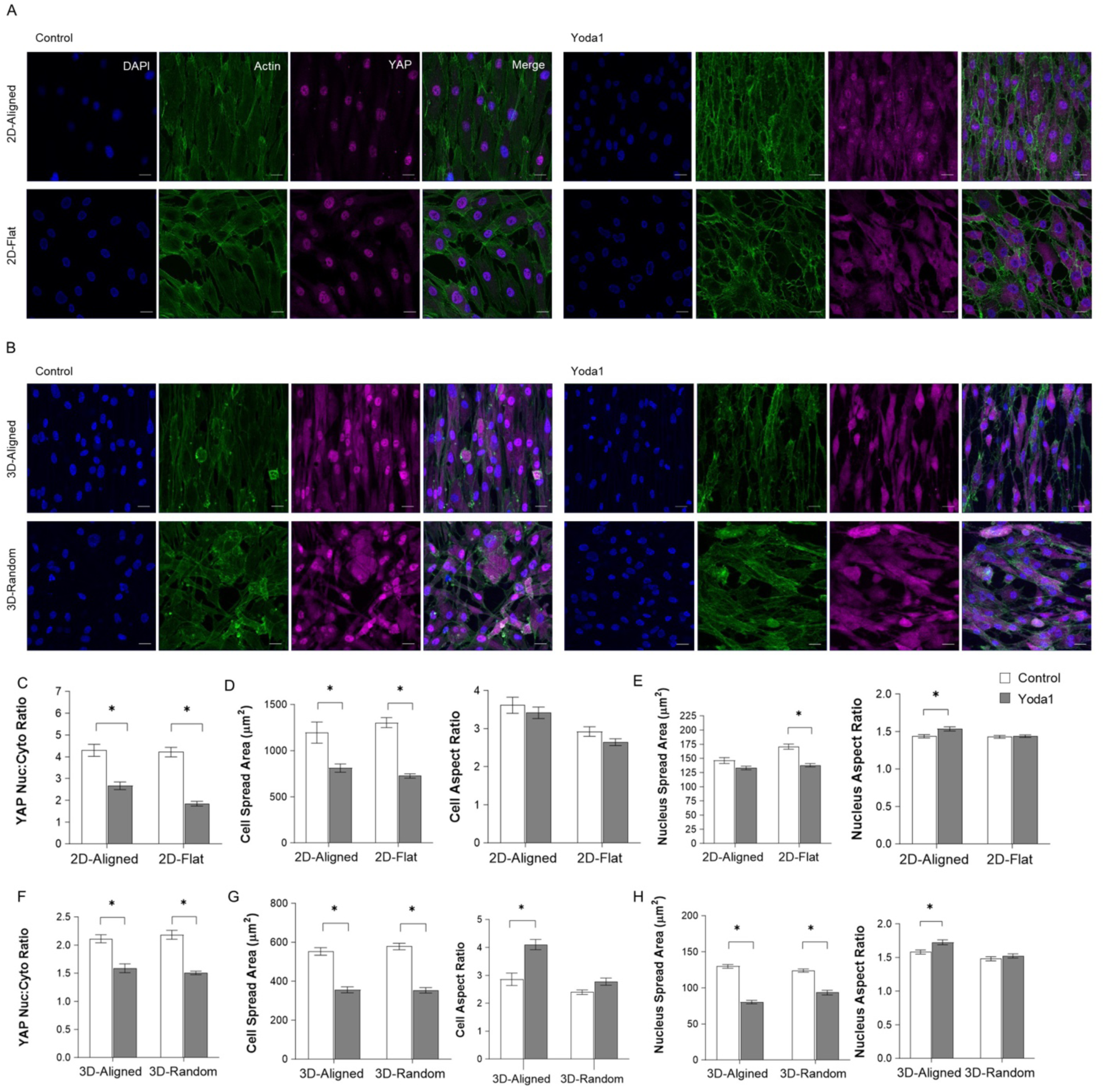
PIEZO1 activation modulates YAP activity and influences cell morphology. Representative nuclear (blue), actin (green), and YAP (pink) staining of control and Yoda1 groups cultured on A) micropatterns (2D model, top) and B) on PCL substrates (3D model, bottom). C) YAP nuclear localization of hASCs cultured on micropatterns (n=3). Quantification of the D) Cell morphology (spread area and aspect ratio) on micropatterns (n=3). E) Nucleus morphology (spread area and aspect ratio) on micropatterns (n=3). F) YAP nuclear localization of hASCs cultured on PCL mats (n=3). Quantification of the G) Cell morphology (spread area and aspect ratio) on PCL mats (n=3). E) Nucleus morphology (spread area and aspect ratio) on PCL mats (n=3). Data presented as mean ± SEM. for group comparison C, D, E, F, G and H, two-way ANOVA with Sidak post hoc test, \**P<*0.05.

In examining the role of PIEZO1 modulation on cell and nucleus morphology, we noted a decrease in hASCs spread area (approximately 30-40 %) following PIEZO1 activation in both 2D and 3D cultures (Fig. 4D, G). This reduction in cell spread area, however, was not accompanied by a significant change in the extent of cell elongation as cell aspect ratio changes with PIEZO1 activation were not significant, except for 3D-aligned cells that we observed cell elongation following PIEZO1 activation (Fig. 4D, G). Furthermore, we observed a statistically significant decrease in nuclear size in cells treated with Yoda1 and cultured on unpatterned, flat surfaces and 3D substrates (Fig. 4E, H). Additionally, in cells grown on aligned substrates, elongation of nuclei was statistically significant both in 2D and 3D (Fig. 4E, H).

### PIEZO1 depletion leads to nuclear accumulation of YAP in hASCs

To validate our observation on the impact of PIEZO1 activation by Yoda1 on YAP localization, we performed *PIEZO1* knock down in cells cultured on both 2D and 3D cell cultures, following the experimental conditions described in the previous section. We expected no significant alteration in YAP localization when *PIEZO1* knocked down ASCs were exposed to Yoda1. Knockdown of *PIEZO1* in hASCs cultured on micropatterns in both aligned and flat conditions resulted in an increased nuclear-to-cytoplasmic (N/C) ratio of YAP (Fig. 5A, C). To discern if this effect was specifically attributable to a loss of PIEZO1 function, we exposed both non-targeting siRNA and *PIEZO1* siRNA groups to Yoda1 and assessed the YAP N/C value as before. Contrary to our expectations, Yoda1 exposure led to a decreased nuclear-to-cytoplasmic YAP ratio in the *PIEZO1* knockdown groups (Fig. 5B, C). Therefore, we hypothesize that the residual PIEZO1 proteins, not eliminated post-siRNA transfection, may still respond to PIEZO1 activation by Yoda1.

**Fig. 5.**
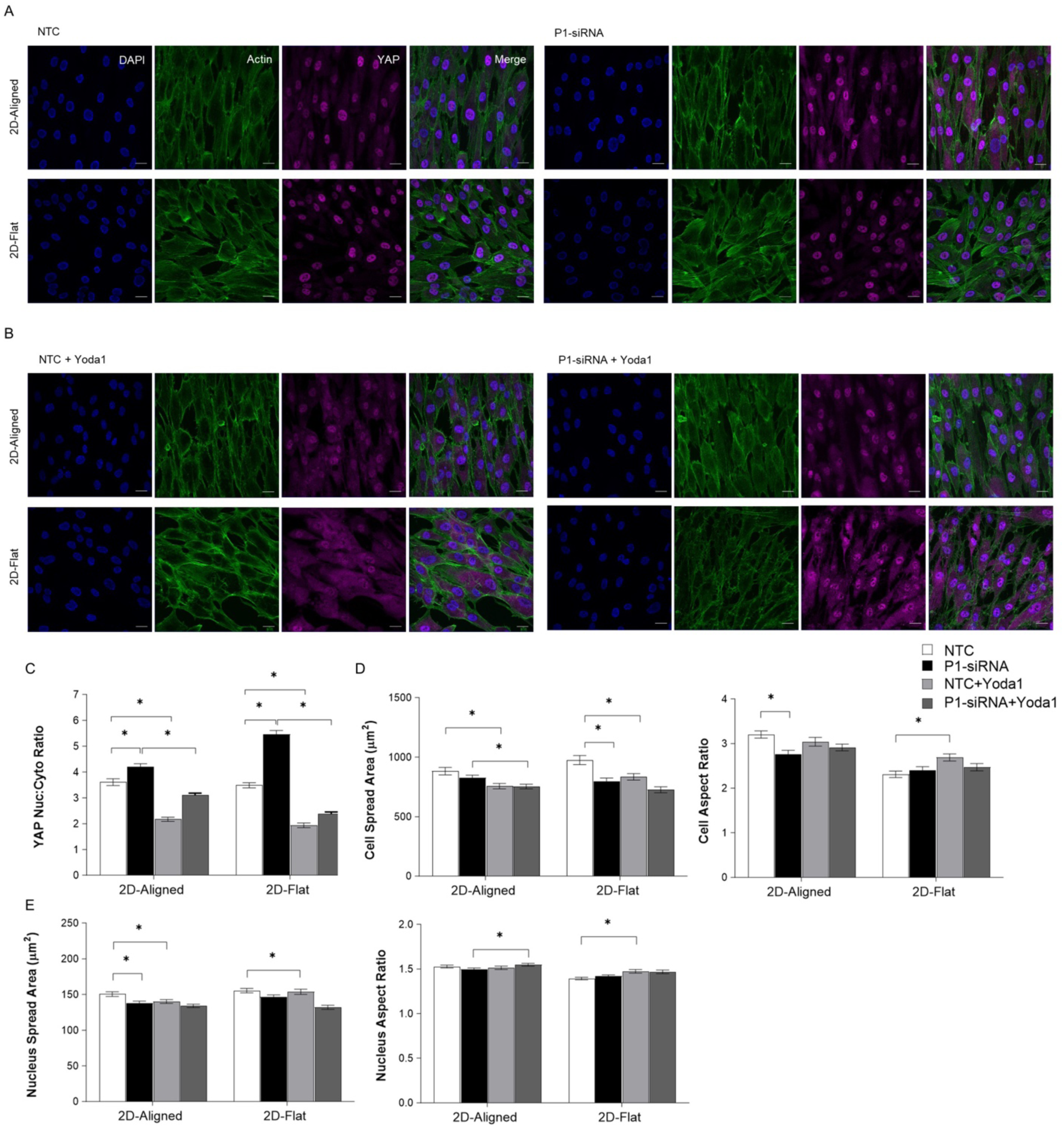
PIEZO1 knockdown modulates YAP activity and influences cell morphology in 2D. Representative nuclear (blue), actin (green), and YAP (pink) staining of A) Non-targeting siRNA and non-targeting siRNA+Yoda1(2.5 μM) groups cultured on micropatterns. B) *PIEZO1* siRNA and *PIEZO1* siRNA+Yoda1 (2.5 μM) groups cultured on micropatterns. Quantification of the C) YAP nuclear localization of hASCs cultured on micropatterns (n=3). Quantification of the D) Cell morphology (spread area and aspect ratio) on micropatterns (n=3). E) Nucleus morphology (spread area and aspect ratio) on micropatterns (n=3). Data presented as mean ± SEM. for group comparison C, D and E, two-way ANOVA with Sidak post hoc test, \**P<*0.05.

Using our 3D model, *PIEZO1* knockdown resulted in an elevated N/C value compared to the non-targeting control group (Fig. 6A, B, C). In the presence of Yoda1, N/C value did not significantly change compared to the *PIEZO1* siRNA groups. Moreover, no significant difference in YAP localization was observed between aligned and random fiber orientations, suggesting that YAP activity in this context is independent of substrate architecture (Fig. 6C).

**Fig. 6.**
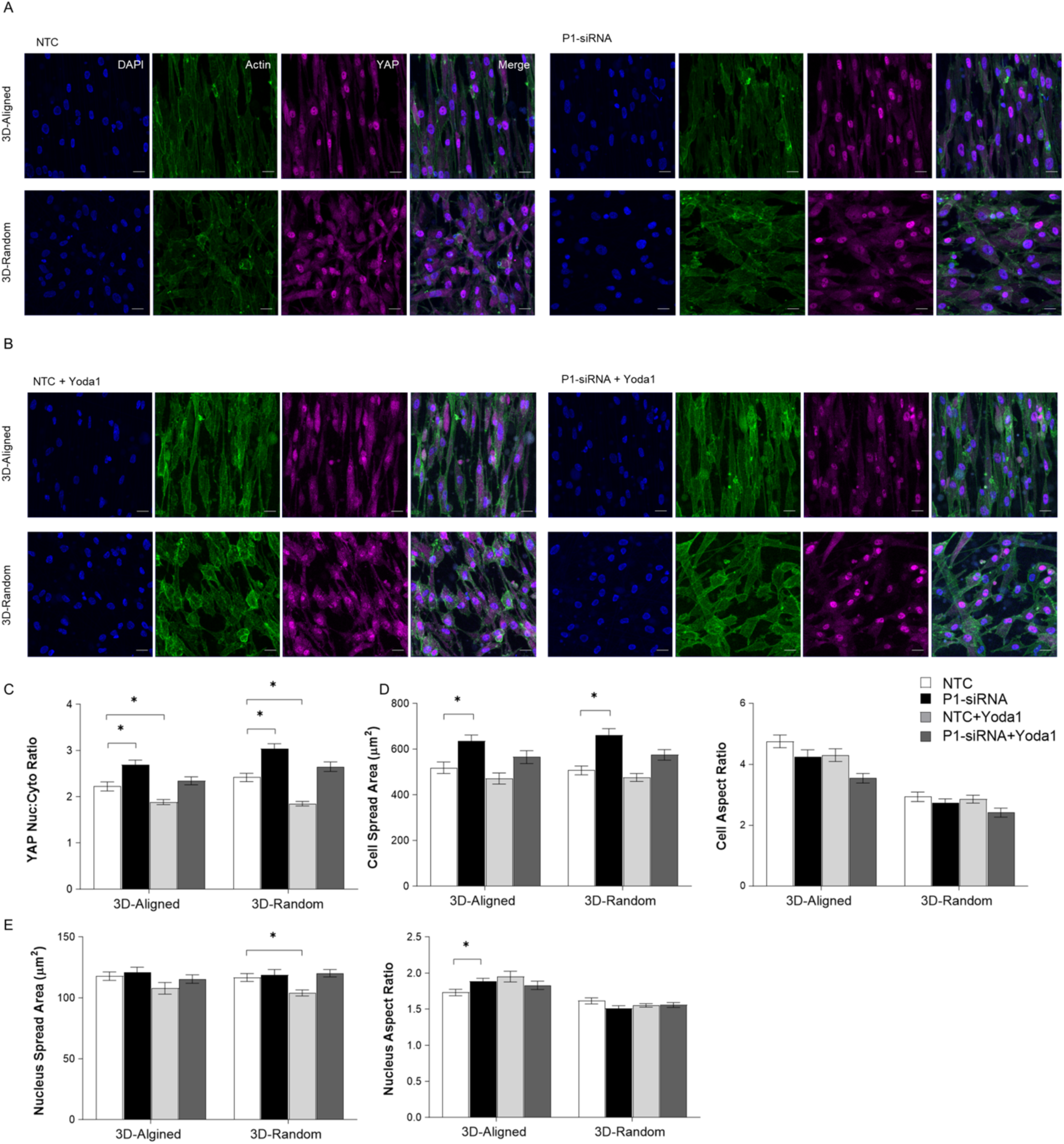
PIEZO1 knockdown modulates YAP activity and influences cell morphology in 2D. Representative nuclear (blue), actin (green), and YAP (pink) staining of A) Non-targeting siRNA, non-targeting siRNA+Yoda1 (2.5 μM) cultured on micropatterns. B) *PIEZO1* siRNA and *PIEZO1* siRNA+Yoda1 (2.5 μM) groups cultured on micropatterns. Quantification of the C) YAP nuclear localization of hASCs cultured on micropatterns (n=3). Quantification of the D) Cell morphology (spread area and aspect ratio) on micropatterns (n=3). E) Nucleus morphology (spread area and aspect ratio) on micropatterns (n=3). Data presented as mean ± SEM. for group comparison C, D and E, two-way ANOVA with Sidak post hoc test, \**P<*0.05.

Our morphological analysis indicated that cultured ASCs only on unpatterned, flat surfaces reduced in size (spread area) following *PIEZO1* knockdown was statistically significant (Fig. 5D). Additionally, hASCs in *PIEZO1* knockdown groups demonstrated reduced elongation compared to their non-targeting siRNA group when cultured on aligned micropatterns (Fig. 5D). The nuclei of hASCs cultured on aligned patterns exhibited reduced spreading when *PIEZO1* was knocked down (Fig. 5E). These results indicate the PIEZO1 contributes to cell spreading and it is involved in cells aligning their morphology in response to matrix alignment.

Expanding our morphological investigation to encompass a 3D model yielded findings that were not paralleled with those observed in the 2D model in the absence of PIEZO1. In the 3D environment, hASCs demonstrated increased spreading on both aligned and random polycaprolactone (PCL) mats (Fig. 6D). Furthermore, an elongation of the nucleus was observed in hASCs cultured on substrates with aligned fibers (Fig 6. E). The specific morphological responses mediated by PIEZO1 appear to be context-dependent, varying between 2D and 3D matrix environments.

### Differential regulation of collagen genes by PIEZO1 activation and YAP/TAZ knockdown in 2D vs 3D models

To elucidate the regulatory role of YAP and TAZ in the context of collagen modulation via PIEZO1, we employed siRNA-mediated gene silencing of the *YAP* and *TAZ* genes. The efficacy of knockdown at both the mRNA and protein levels was evaluated through RT-qPCR and Western blot analyses. The results demonstrate a successful reduction in *YAP* and *TAZ* expression (Fig. 7A, B).

**Fig. 7.**
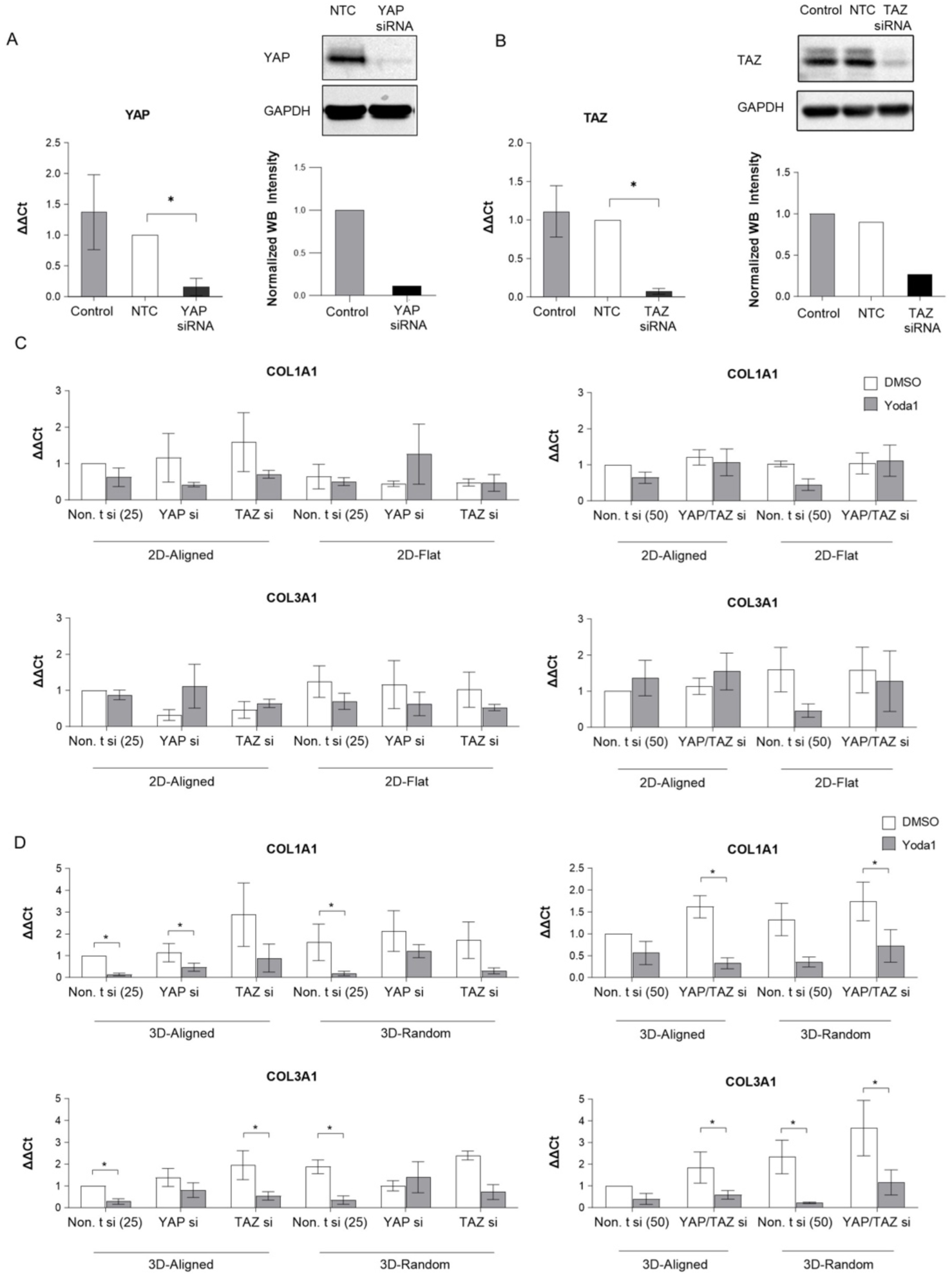
PIEZO1 mechanotransduction in collagen regulation of hASCs is not dependent on YAP and TAZ. A) YAP is efficiently knocked down in mRNA and protein levels. mRNA levels of YAP normalized to *GAPDH* expression level in non-targeting siRNA and YAP siRNA groups (left). Protein levels of non-targeting siRNA and *YAP* siRNA groups and their normalized intensity values (right) from western blotting. B) *TAZ* is efficiently knocked down in mRNA and protein levels. mRNA levels of *TAZ* normalized to GAPDH expression level in non-targeting siRNA and *TAZ* siRNA groups (left). Protein levels of non-targeting siRNA and *TAZ* siRNA groups and their normalized intensity values (right) from western blotting. C) mRNA levels of *COL1A1* normalized to *GAPDH* in response to *YAP* siRNA+Yoda1, *TAZ* siRNA+Yoda1, and *YAP/TAZ*+Yoda1 compared to the same siRNA groups without Yoda1 exposure on Micropatterns and PCL mats. D) mRNA levels of *COL1A1* normalized to *GAPDH* in response to *YAP* siRNA+Yoda1, *TAZ* siRNA+Yoda1, and *YAP/TAZ*+Yoda1 compared to the same siRNA groups without Yoda1 exposure on Micropatterns and PCL mats. Data presented as mean ± SEM. for group comparison A and B, one-way ANOVA with Dunn post hoc test, \**P<*0.05, for C and D, multiple t-test, \**P<*0.05.

To investigate the impact of YAP and TAZ depletion, individually and in combination, on collagen gene expression during PIEZO1 activation, we used RT-PCR to compare the collagen gene expression patterns of cells treated with or without Yoda1 exposure, in both 2D and 3D substrate contexts (Fig. 7C, D). As we previously observed, PIEZO1 activation with Yoda1 did not have significant effect on collagen gene expressions (Fig. 2E), we also found here that activation of PIEZO1 in the context of *YAP* or *TAZ* knockdown, as well as the concurrent knockdown of both *YAP* and *TAZ*, does not significantly alter collagen expression patterns on micropatterned substrates (Fig. 7C). *COL1A1* gene expression was significantly downregulated in both aligned and random substrate cultures in 3D culture when hASCs with *YAP/TAZ* knockdown and *YAP* knockdown cells were exposed to Yoda1 (Fig. 7D).

In the 3D model, a reduction in COL3A1 expression was observed in both aligned and random substrate cultures within the *TAZ* and combined *YAP/TAZ* siRNA groups (Fig. 7D). The effect of PIEZO1 activation on collagen gene expressions in the absence of YAP and/or TAZ was more pronounced in a 3D model as compared to a 2D model.

## Discussion

In this study, we identified a role for PIEZO1 and the mechano-responsive transcriptional activators YAP/TAZ in sensing substrate architecture and regulating collagen production in human ASCs (Fig. 8). Using both 2D and 3D nanostructured in-vitro models, we explored the interaction between PIEZO1 and the YAP/TAZ signaling pathway in this process. Our findings demonstrated that PIEZO1 activity can influence collagen expression and morphological responses of hASCs in a manner dependent on the substrate architecture. Interestingly, PIEZO1 knockdown led to increased collagen production, suggesting a regulatory role in limiting collagen synthesis. This PIEZO1-mediated collagen modulation was sensitive to the substrate architecture, with a significant effect observed when hASCs were cultured on aligned substrates in both 2D and 3D contexts. Our 3D model revealed reduced collagen I protein synthesis upon PIEZO1 activation. Furthermore, we observed decreased nuclear accumulation of YAP by PIEZO1 activation by Yoda1, indicating that PIEZO1 activity influences YAP localization. While YAP activity was influenced by PIEZO1 activity, our gene expression analysis did not reveal a dependency of YAP/TAZ on PIEZO1 in the regulation of collagens in these settings.

**Figure 8.**
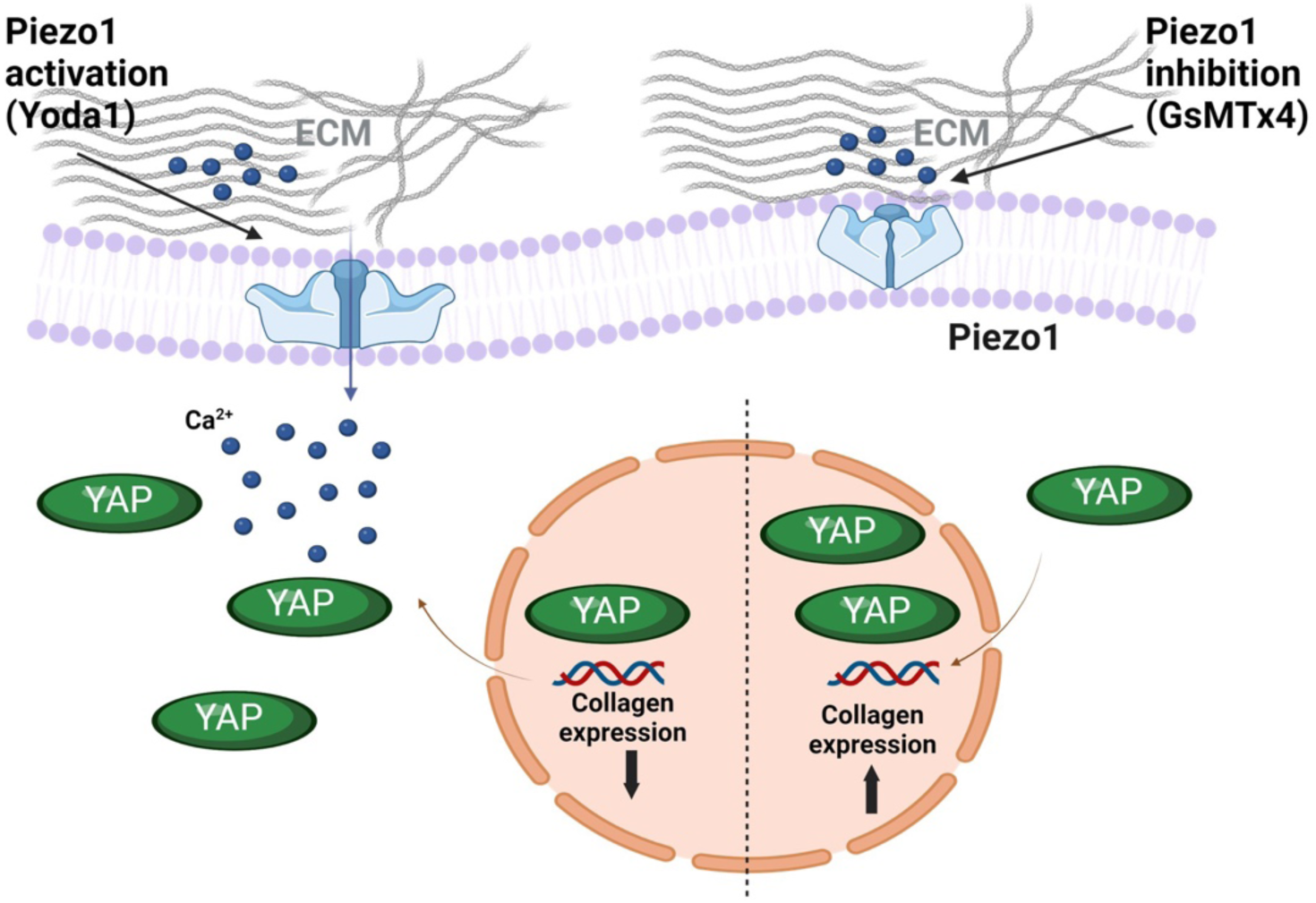
Regulation of collagen expression in hASCs by PIEZO1 modulation via YAP/TAZ dependent mechanotransduction. Schematic illustrating Regulation of collagen expression in ASCs by PIEZO1-mediated mechanotransduction and PIEZO1 interaction with YAP.

Our findings contrast with a recent study proposing PIEZO1 inhibition as a potential strategy for mitigating fibrosis, particularly in organs where adipose tissue contributes to fibrotic processes [31]. Intriguingly, our data reveal an opposing effect, wherein inhibition or knock-down of PIEZO1 led to increased collagen production in both 2D and 3D microenvironments using human adipose-derived stem cells. This result suggests that PIEZO1 may function as a negative regulator of collagen synthesis in hASCs, exerting anti-fibrotic effects. These observations challenge the proposed therapeutic benefit of PIEZO1 inhibition and instead implicate PIEZO1 as a potential target for promoting the resolution of fibrosis in adipose tissue. The contrasting role of PIEZO1 in regulating collagen production and potentially fibrosis using ASCs compared to other tissues or cell types can be reconciled by considering that PIEZO1 may exhibit differential effects on collagen synthesis depending on the cell-type specific functions; it may be different in ASCs compared to fibroblasts or myofibroblasts, which are more directly involved in fibrosis progression. Moreover, microenvironmental cues and mechanical forces experienced by cells in different tissues can influence the downstream signaling pathways and cellular responses mediated by PIEZO1. PIEZO1 may play divergent roles in regulating collagen production and matrix remodeling during different developmental stages, and functional contexts. For instance, its effects on hASCs, which contribute to adipose tissue homeostasis and regeneration, may differ from its role in chronic disease states or injury-induced fibrotic processes in other tissues. Additionally, Differences in experimental conditions, such as the use of 2D or 3D culture models, the specific extracellular matrix components or substrates utilized, or the duration and extent of PIEZO1 modulation, could contribute to the observed contrasting effects on collagen regulation.

The greater impact of *PIEZO1* knockdown on collagen expression in hASCs grown on aligned as opposed to random substrates, may emphasize the significance of substrate architecture on cellular behavior and PIEZO1 mechanotransduction pathways. This aligns with evidence showing substrate fiber alignment impact on cell behavior, differentiation, gene expression, and cytoskeletal organization [58-63]. For instance, in a study Kishore et al. [64] demonstrated that aligned collagen fibers more effectively promote the expression of tendon-specific markers, including collagen III, when compared to randomly oriented fibers. They concluded that aligned ECM fibers possess an enhanced capacity to drive the differentiation of human mesenchymal stem cells towards a tendon-like phenotype. This highlights the significant impact of substrate architecture on regulating gene expression specifically collagens.

Furthermore, PIEZO1 activation induced significant morphological changes by reducing cell spreading area and increasing cell elongation in hASCs, especially in cells cultured on aligned substrates, ASCs elongation was significant linking PIEZO1 mechanotransduction to regulation of cell morphology. This is in line with existing literature highlighting the role of PIEZO1 in cellular morphology changes [56, 65-70]. Holt et al. [65] explored PIEZO1 localization in migrating keratinocytes during wound healing. Their results indicate that PIEZO1 activity regulates cell shape by promoting cell polarization in keratinocytes, potentially by modulating the subcellular localization of PIEZO1 channels in migrating cells. In another study on HEK cells cultured on stripe patterns, Jetta et al [56] showed that PIEZO1 regulates cell spreading. PIEZO1-expressing HEK cells exhibited elongated morphology, which was abolished by *PIEZO1* knockout as well as inhibition with GsMTx4 [56]. This insight is particularly relevant for understanding cell responses to their mechanical environment, crucial in tissue engineering. Our results also revealed that PIEZO1 activation induces YAP accumulation in the cytoplasm, resulting in reduced YAP activity. Multiple studies have shown interactions between these pathways and their functional implications in cellular processes in other tissues/cells [41, 71-75]. Xiong et al. [71] demonstrated that PIEZO1 activation promotes ovarian cancer metastasis via the Hippo/YAP signaling axis. The Hippo pathway, which includes the downstream effectors YAP and TAZ, is a critical regulator of organ size control and stem cell renewal [76, 77]. Given the involvement of the Hippo pathway in organ size control and stem cell renewal, these findings suggest that PIEZO1 could potentially influence cellular processes related to tissue homeostasis and regeneration through its effects on the Hippo/YAP signaling cascade. Recent research findings have indicated that the absence of PIEZO1 causes YAP to be excluded from the nucleus in human neural stem cells [41] and during zebrafish outflow tract valve development [73], implying that PIEZO1 may function upstream of YAP. PIEZO1 and PIEZO2 have also been identified as upstream regulators of signal transduction pathways that activate YAP for stem cell differentiation [74], also in regulating self-renewal of vocal fold mucosal epithelia [75].

However, in another study, it was demonstrated that YAP signaling induces PIEZO1 to promote oral squamous cell carcinoma cell proliferation [72]. Interestingly, knocking down PIEZO1 did not influence the transcriptional targets of YAP or YAP/TAZ nuclear localization. Consequently, the authors propose that while the interplay between YAP signaling and PIEZO1 may vary depending on the cellular environment, these molecules could serve as a shared mechanotransduction mechanism, bridging organ development in stem cells [72].

Our findings suggest that YAP and TAZ do not significantly influence collagen gene expression when PIEZO1 is activated on micropatterned substrates. This indicates independence between the mechanotransduction pathway mediated by PIEZO1 and the signaling mediated by YAP/TAZ in regulation of collagens. These results contrast with some previous research on the interplay between YAP/TAZ and PIEZO1 in collagen expression in other cell types. For example, Wang et al. [78] studied osteoblast cells response to mechanical loads and mechanistically investigated expression of collagens. Their findings suggested that PIEZO1 regulated the expression of type II and IX collagens through YAP. Another study on heart valve interstitial cells suggested that PIEZO1 activation triggered Ca^2+^ influx and YAP translocation to the nucleus, further supporting the role of YAP in PIEZO1-mediated collagen regulation [68]. Our findings suggest that additional molecular pathways need to be investigated to gain a better understanding of the downstream mechanisms involved in this process.

Finally, this study unveiled a previously unexplored role for the PIEZO1 ion channel and its interplay with the YAP/TAZ signaling pathway in regulating collagen production and morphological responses of human adipose-derived stem cells (hASCs) to substrate architecture. Utilizing both 2D and 3D nanostructured in vitro models, we demonstrated that PIEZO1 activity modulates collagen expression and cellular morphology in hASCs, with a more pronounced effect on aligned substrates. Interestingly, PIEZO1 depletion led to increased collagen production, suggesting a regulatory role in limiting collagen synthesis. Furthermore, PIEZO1 activation induced YAP cytoplasmic accumulation, indicating its influence on YAP localization. However, gene expression analysis revealed no dependency of YAP/TAZ on PIEZO1 in regulating collagens in these settings, suggesting potential independence between these pathways in hASCs. Our findings suggest that additional molecular pathways need to be investigated to gain a better understanding of the downstream mechanisms involved in this process. In future studies, exploring the interplay between PIEZO1 and other mechanotransduction pathways, such as integrin signaling, will be crucial in understanding how cells respond to substrate topography and mechanical cues to regulate collagens in disease context. Future studies may wish to investigate potential crosstalk and feedback mechanisms between PIEZO1 and YAP/TAZ signaling pathways across various cellular contexts and developmental stages. Previous research indicates a bidirectional relationship between PIEZO1 and YAP/TAZ signaling, underscoring the importance of gaining deeper insights into their interplay.

## Acknowledgements

This work was supported by the Shriners Hospitals for Children and the National Institutes of Health (AG15768, AG46927, AR080902, AR072999, AR073752, AR074992). We thank the Institute of Materials Science and Engineering (IMSE) at Washington University in St. Louis for their support and training on fabrication of micropatterned substrates. We thank Dr. Elizabeth Haswell for her insights and editing of the manuscript.

## Data Availability Statement

The raw/processed data required to reproduce these findings cannot be shared at this time due to technical or time limitations.

