## Supplemental Information for "PIEZO1-Mediated Mechanotransduction Regulates Collagen Synthesis on Nanostructured 2D and 3D Models of Fibrosis"

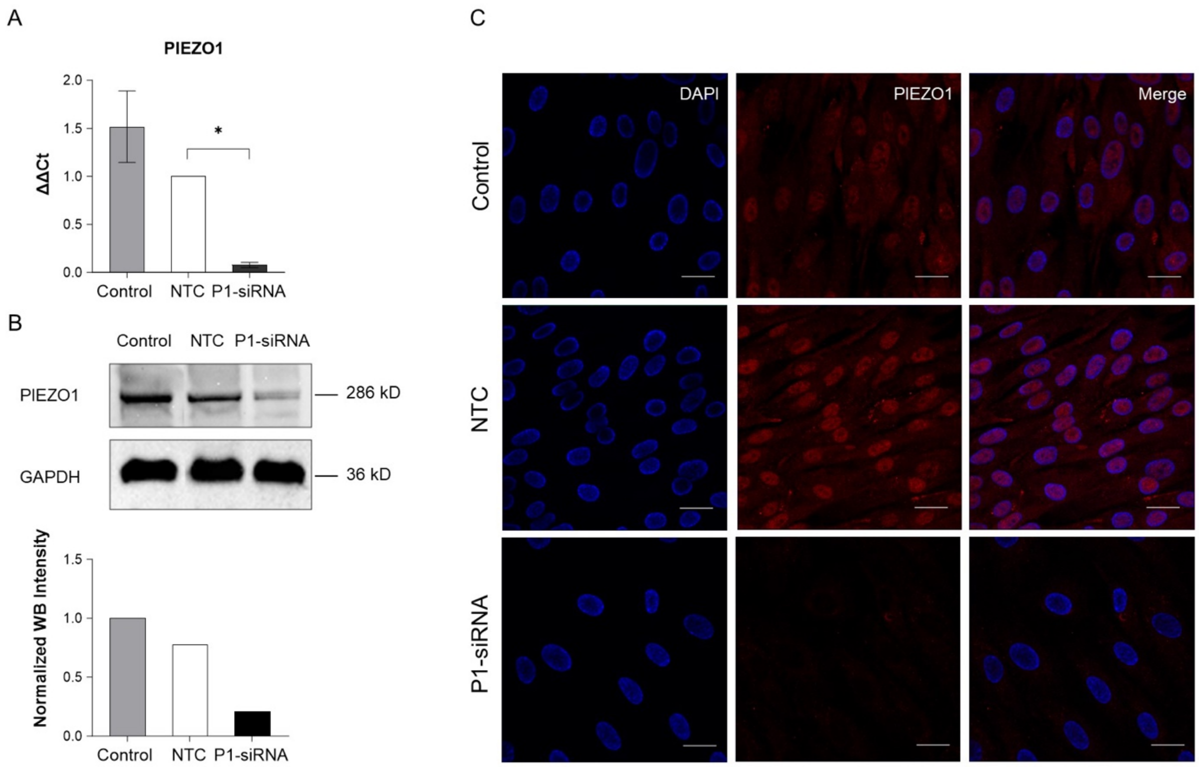


**Supplementary Fig. S1.** A) mRNA levels of PIEZO1 normalized to GAPDH expression level in control (transfection reagent, no siRNA), non-targeting (NT) siRNA, and PIEZO1 siRNA hASCs. B) PIEZO1 protein expression in control (transfection reagent, no siRNA), NT siRNA, and PIEZO1 knock-down hASCs, and normalized intensity values of protein expression quantified using WB (left). C) Protein expression by IF. Representative nuclear (blue) and PIEZO1 (red) staining of hASCs cultured on cover glass for control, transfected with NT siRNA and PIEZO1 siRNA (Right), scale bar: 20 µm. Data presented as mean ± SEM. for E, n=4 samples, $^{*}P<0.05$

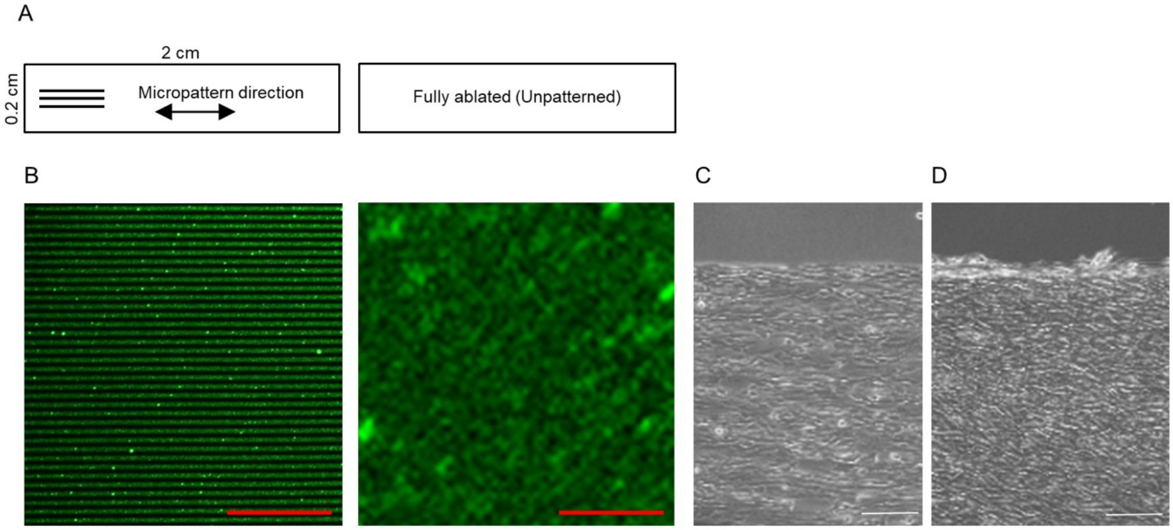


**Supplementary Fig. S2.** A) Schematic of micropattern design (left: aligned micropattern, right: unpatterned). B) Fibronectin adsorption on microphotopatterned substrates. Left: 2 µm wide patterns with 5 µm center-to-center distance. Right: Fully- ablated pattern area. Scale bar is 50 µm. C) Brightfield images of hASCs on micropatterns after an overnight post-seeding and D) after 7 days. Scale bar is 200 µm.

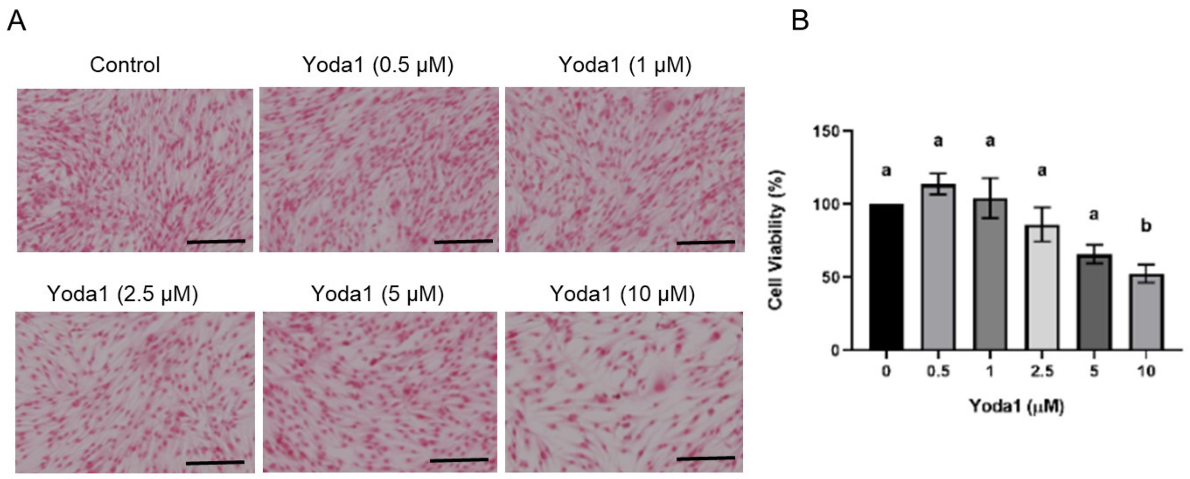


**Supplementary Fig. S3.** A) Representative brightfield images of hASCs stimulated with different daily doses of Yoda1. Scale bar is 200 µm. B) hASC viability in response to Yoda1 treatment assessed by MTT assay, n=3. One-Way ANOVA, different letters indicate statistically significant results, $p<0.05$.

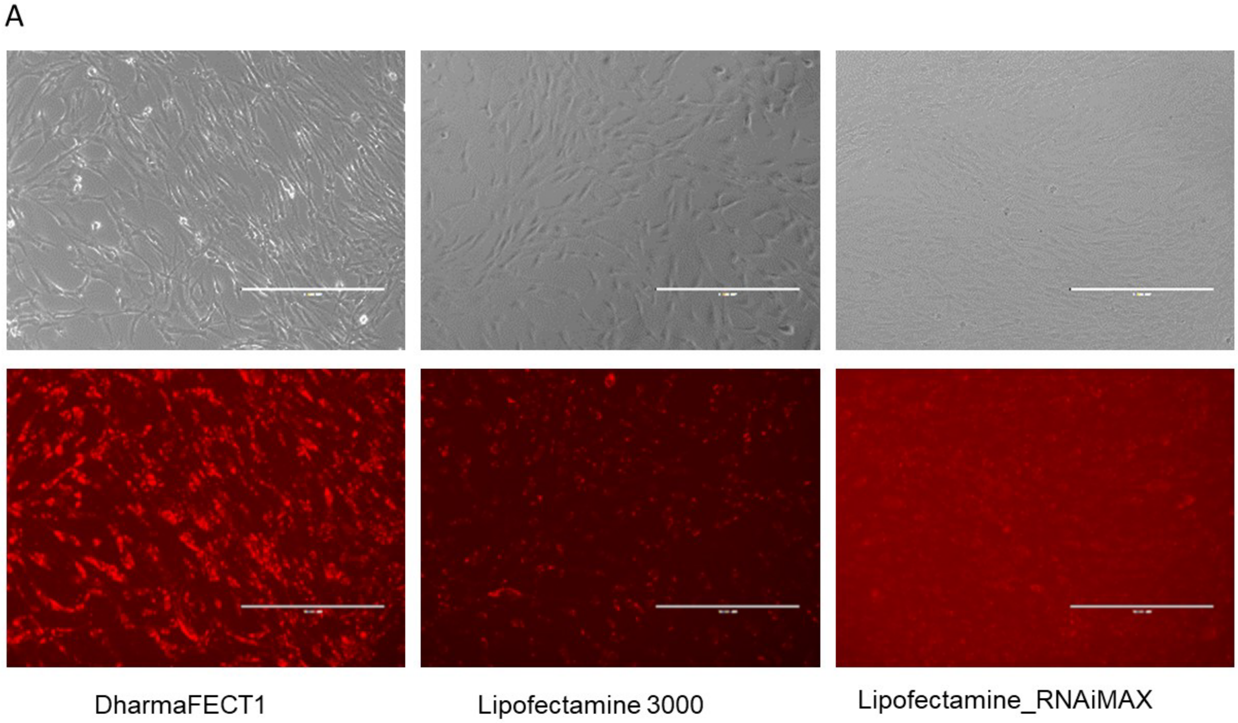


**Supplementary Fig. S4.** A) Optimization of efficient siRNA delivery using fluorescent siGLO Lamin A/C Control siRNA (ed) and different transfection reagents. Scale bar is 1000 µm.

**Supplementary Table. 1.** Real time PCR primer sequences.

| Gene | Primer-forward | Primer-Reverse |
|---|---|---|
| GAPDH | cagcctcaagatcatcagca | tgtggtcatgagtccttcca |
| COLLAGEN I | gtgctaaaggtgccaatggt | accaggttcaccgctgttac |
| COLLAGEN III | ccaggagctaacggtctcag | cagggtttccatctcttcca |
| PIEZO1 | tggcaacaaaatcagcagag | gccattgtcaacagcagaga |
| YAP | cccaactggcattgactttt | ctcgaggtctcccactgaag |
| TAZ | gtcaccgtgtcaacctgatg | gcctgccttcaagatttctg |

**Supplementary Table. 2.** Sequence of siRNA pools.

| Gene | siRNA pool sequences |
|---|---|
| GAPDH | GUCAACGGAUUUGGUCGUA, CAACGGAUUUGGUCGUAUU, GACCUCAACUACAUGGUUU, UGGUUUACAUGUUCCAAUU |
| Non-targeting | UGGUUUACAUGUCGACUAA, UGGUUUACAUGUUGUGUGA, UGGUUUACAUGUUUUCUGA, UGGUUUACAUGUUUUCCUA |
| PIEZO1 | GCAGCAUGACAGACGACAU, UGGGUAUGCCAACGAGAA, UGGCUGAUGUUGUCGACUU, GCGCAUCAGUCAUCGUUUU |
| YAP | GCACCUAUCACUCUCGAGA, UGAGAACAAUGACGACCAA, GGUCAGAGAUACUUCUUAA, CCACCAAGCUAGAUAAAGA |
| TAZ | CCACCAAGCUAGAUAAAGA, GGACAAACACCCAUGAACA, AGGAACAAACGUUGACUUA, CCAAAUCUCGUGAUGAAUC |
